# Simultaneous visualization of cells and marker genes from scRNA-seq studies

**DOI:** 10.1101/2022.12.27.521966

**Authors:** Namrata Bhattacharya, Swagatam Chakraborti, Krishan Gupta, Aayushi Mittal, Debajyoti Sinha, Colleen Nelson, Tanmoy Chakraborty, Gaurav Ahuja, Debarka Sengupta

## Abstract

The complexity of scRNA-sequencing datasets highlights the urgent need for enhanced clustering and visualization methods. Here, we propose Stardust, an iterative, force-directed graph layouting algorithm that enables the simultaneous embedding of cells and marker genes. Stardust, for the first time, allows a single-stop visualization of cells and marker genes on a single 2D map. While Stardust provides its own visualization pipeline, it can be plugged in with state-of-the-art methods such as Uniform Manifold Approximation and Projection (UMAP) and t-Distributed Stochastic Neighbor Embedding (tSNE). We benchmarked Stardust against popular visualization and clustering tools on both scRNA-seq and spatial transcriptomics datasets. In all cases, Stardust performs competitively in identifying and visualizing cell types in an accurate and spatially coherent manner.

## INTRODUCTION

Messenger RNAs (mRNAs) are positioned at the center of genetic information flow and cast phenotypic dynamicity to a cell to help it adapt to its environment and external impulses. Microfluidic breakthroughs over the past few years have enabled high-throughput measurement of mRNAs at the single cell resolution [1]. The recent emergence of spatial transcriptomics techniques has added a much-needed dimension to the field by enabling spatially resolved characterization of cellular diversity *in situ*, within a tissue [2,3]. Further to it, today, multiplexed single-molecule Fluorescence in situ hybridization (FISH) measurements allow studying subcellular compartmentalization of RNA molecules [4]. Together, these breakthroughs have provided several degrees of freedom to our interrogation of complex biological systems, under normal, developmental or pathological conditions. Harnessing such a multi-factorial molecular information deluge requires the assistance of unsupervised learning methods. As such, efficient dimension reduction and clustering of single cell expression profiles play a pivotal role in the systemic appraisal of cell-to-cell variability [5,6]. Remarkably, the progressive evolution of such computational methods underscores the yet unsettled nature of this field [7–9].

Single cell RNA sequencing (scRNA-seq) data are inherently noisy due to the small amounts of starting RNA material and cell-to-cell technical biases [10,11]. Trivial biological variability due to cell cycle stage, response to pathogens, cell viability, differential replicative age, etc., further impedes expression-based characterization of cellular heterogeneity. While the advent of novel clustering methods is significantly fast-paced, dimension reduction techniques transcend slowly. Recently, the field has embraced Uniform Manifold Approximation and Projection (UMAP) as the best practice method for the low-dimensional embedding of single cells [9] due to improved segregation of distinct cell-types as compared to t-Distributed Stochastic Neighbor Embedding (t-SNE), and its ‘linear’ ancestor Principal Component Analysis (PCA). UMAP’s wide adoption highlighted the importance of spatial segregation of transcriptomically distinct cell-types in a 2D portrait of cells. Despite significant improvements, cell-clusters with remarkably different lineages are found on a transitional continuum of cellular identity with loosely defined boundaries. This frequently contradicts with clustering outcomes. We ask ourselves - “How do cells form distinct clusters yet run into each other’s territory on a low-dimensional map?”. This leads to the conclusion that the existing dimensionality reduction techniques might not be able to manifest transcriptional heterogeneity adequately, as otherwise teased out by the clustering algorithms. Due to this inability, abstract dimensions obtained from feature extraction/dimension reduction techniques are usually not used as features for clustering. Another practical difficulty is cross-referencing between the 2D map and the differential expression heatmap. For instance, the recently published Human Cell Landscape (HCL) reported 102 clusters obtained from about ∼700,000 cells from ∼50 human tissues [12]. The sheer number of potential cell-types creates cognitive overload while inspecting clusters and their marker genes across the low-dimensional visualization and the differential expression heatmap. To address these challenges, we implemented Stardust, an iterative graph-layouting algorithm for cell-gene co-embedding that teases out transcriptional heterogeneity in a manner that preserves cell-lineage identities without getting into the trap of over-clustering.

## RESULTS AND DISCUSSION

### Overview of Stardust

Stardust integrates genes and single cells as part of the same network topology, with two legal categories of connections/edges: cell-cell and cell-gene based on nearest neighbor principles. While cell-cell connections are established based on their expression similarity, a gene is connected to a few top cells, wherein its relative abundance is high. On the first pass, genes are selected using Principal Component Analysis (PCA). For graph embedding, Stardust uses OpenOrd [13], a scalable and qualitatively improved variant of the force-directed graph layouting algorithm. Cell-clusters obtained from OpenOrd-based low-dimensional embedding are subjected to Differential Expression (DE) analysis. DE genes thus obtained replace the PCA selected genes, which change the network conformation and thus the cell clusters. Clustering, DE analysis, and OpenOrd based layouts are performed iteratively until we arrive at stable clusters. Typically this is achieved in 3 iterations. In the end i.e. on the 4^th^ pass, for a given expression matrix, Stardust outputs 2D embedding of cells/DE genes, and their inferred cluster identities. Cell-type specific genes get naturally inscribed within the associated cell-clusters, thereby simplifying the visual interpretation of tissue heterogeneity. Besides OpenOrd based embeddings, due to the wide acceptance of UMAP, Stardust also allows the user to visualize clusters and the associated marker genes within a UMAP defined 2D space (**Supplementary Figure S1**).

### Comparative benchmarking of Stardust visualization and clustering

We benchmarked Stardust visualization and clustering against numerous best practice methods. Notably, Stardust enables cell-gene co-embedding in a unified 2D space, which is compatible with all dimension reduction tools. In particular, we evaluated UMAP [14], t-SNE [15], and SPRING [16] for visualization and Louvain [17], Leiden [17], SIMLR [18], and SC3 [19] for clustering, due to their wide acceptance in the community. For performance assessment, we acquired six publicly available scRNA seq datasets, of which four had cell type annotation available at the single cell level. We applied Stardust UMAP, tSNE and SPRING to demonstrate accurate placement of DE genes across 2D visualization of given single cells (**Figure 1a-d**). In each visualization, clusters are obtained using Stardust. For each cluster, a representative upregulated gene (the top one) can be found inscribed within the respective clusters. To avoid overcrowding of the plots, Stardust empowers the user to choose the genes to annotate. Notably, we found Stardust visualization is highly consonant with UMAP based 2D organization of the cells. To assess the quality of clusters, we considered four datasets *PBMC, Melanoma, Pollen* and *Kolod*. In each case, we benchmarked the clustering outcomes against cell annotations from an external source. We recorded normalized mutual information (NMI) for Stardust alongside widely used clustering methods Louvain (from scanpy), Leiden (from scanpy), SIMLR and SC3. Stardust performed the best in terms of the recorded average NMI value. Whereas, Louvain ranked second.

**Figure 1.**
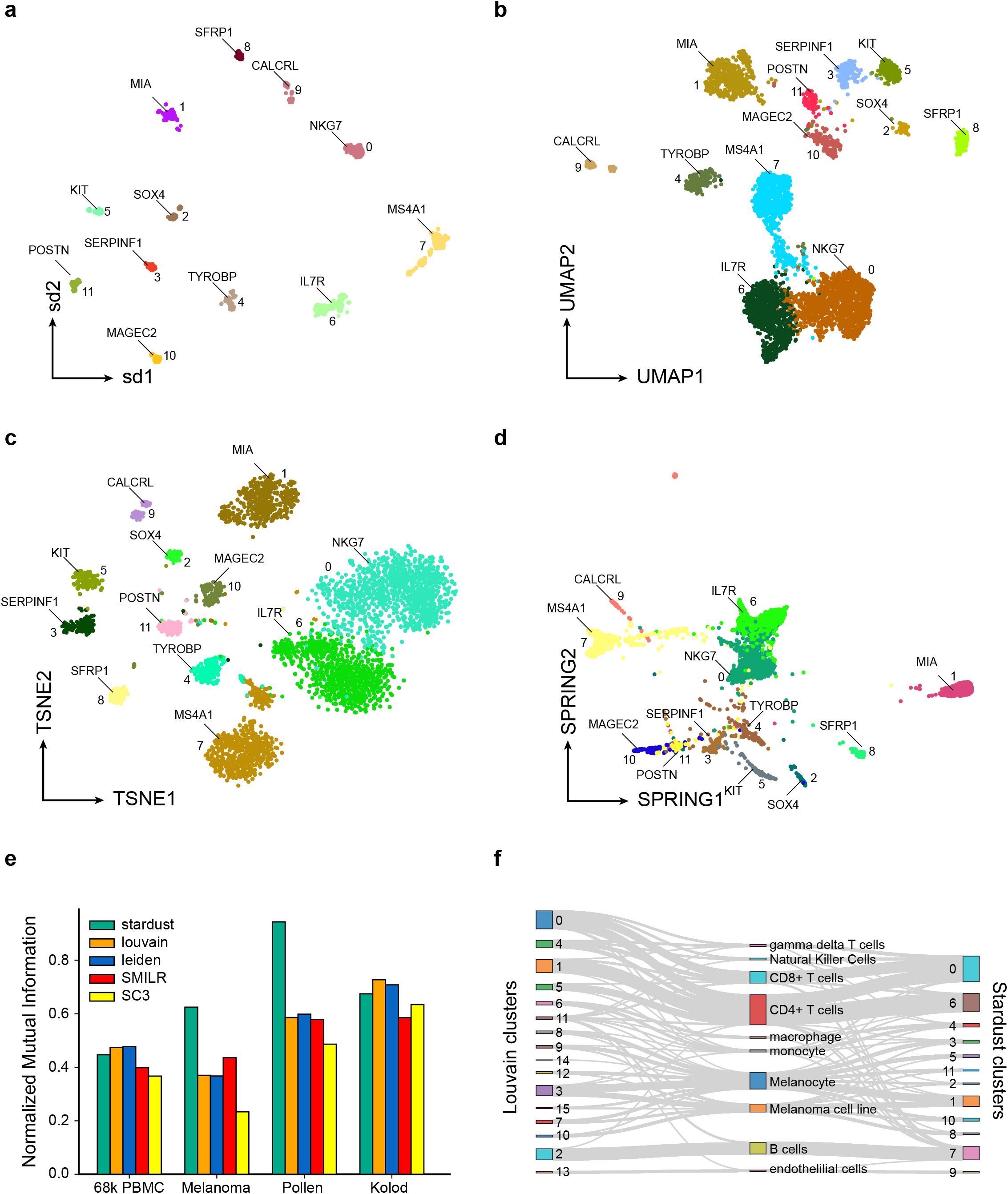
Stardust based visualization and clustering of scRNA-seq profiles. (a) UMAP based visualization of the *Melanoma* data, with marker gene locations inferred by Stardust. (b) Stardust based co-embedding of cells and genes from the *Melanoma* dataset. (c) Alluvial plot depicting the correspondence between unsupervised clusters obtained using both the methods, and marker-based annotation of the single cells using scMatch. (d) Normalized Mutual Information (NMI) computed on *PBMC* and *Melanoma* datasets cross-referencing the clusters with cell annotations. (e) Silhouette index computed on all the datasets to measure signal-to-noise ratio associated with 2D locations of the cell clusters.

We critically examined clusters obtained by Stardust and Louvain on the *Melanoma* data. *Melanoma* data comprises 4645 scRNA-seq profiles from 19 patients, spanning both malignant and non-malignant cell-subtypes from human melanoma tumors. Marker-based bioinformatic annotations of the cells were obtained from the scMatch R package [20]. scMatch mapped the malignant cells into two categories based on its reference database i.e. melanoma cell lines and primary melanocytes. In the case of Stardust, besides T cell and monocyte subtypes, the majority of the nonmalignant cells were concentrated into a single cluster. As expected, the malignant cells were found distributed across multiple clusters, underscoring inter/intratumoral heterogeneity harbored by the malignant cells (**Supplementary Figure S2a**). Conversely, Louvain-based clustering using scanpy fragmented rather homogeneous populations of CD4+ and CD8+ cell-types into multiple clusters giving rise to discordance with annotations (**Supplementary Figure S2b**). A major cell-type segregation ambiguity was spotted in cluster 4 (resulted by Louvain clustering), which shared a considerable number of cells from three scMatch annotated subpopulations namely CD4+/CD8+ T cells and Natural Killers. All these results collectively suggest that Stardust outperforms the current state of the art both in low-dimensional visualization of cells and clustering.

We repeated Stardust clustering and visualization using multiple additional datasets — PBMC data (∼68,000 annotated cells), Liver data (∼7000 cells with bioinformatically inferred cell types) [21], mouse brain data (∼20000 cells with bioinformatically inferred cell types) [22]. In all cases we found Stardust visualization, clustering, and marker gene co-embedding to be highly consonant with standard methods, author annotation and ground truth annotations, where available (**Supplementary Figures S3-S5**). For all datasets except *Kolod*, and *Pollen*, where the number of cells is less, we computed and reported Silhouette scores of the clusters using Stardust and Louvain. Stardust consistently outperformed Louvain on all datasets (**Supplementary Figure S6**). Cluster-specific top differentially upregulated genes obtained using Stardust and scanpy (Louvain for clustering) are tabulated in a study-specific manner (**Supplementary Table 1**).

### Stardust accurately segregates influenza virus infected cells

In the above comparative studies, we observed that Stardust visualization offers a higher signal-to-noise ratio for cell-type detection and segregation. To assess it further, we posed an independent and stricter challenge. Sun and colleagues profiled about 9000 cells, including uninfected A549 cells (originating from lung) as control, exposed to H1N1 but uninfected cells (bystanders) and exposed and infected cells [23]. The distinction between bystanders and infected cells was defined bioinformatically based on the availability of viral RNA traces in the single cells. We hypothesized that cells with viral nucleic acid traces would have a different phenotype as compared to the bystanders. Visual separation of these infected cells thereby serves as a criterion to assess the performance of various dimension reduction techniques. To this end we subjected the single cells to visualization using Stardust, alongside other widely used methods — UMAP, tSNE and SPRING. Notably, SPRING is comparable to Stardust since it is based on networks. All methods performed reasonably well in visually segregating the infected cells (**Figure 2a-d**). Leakage of infected cells to other clusters or co-clustering of uninfected/bystander cells with the infected cells could be due to imperfections associated with bioinformatic annotation of cells.

**Figure 2.**
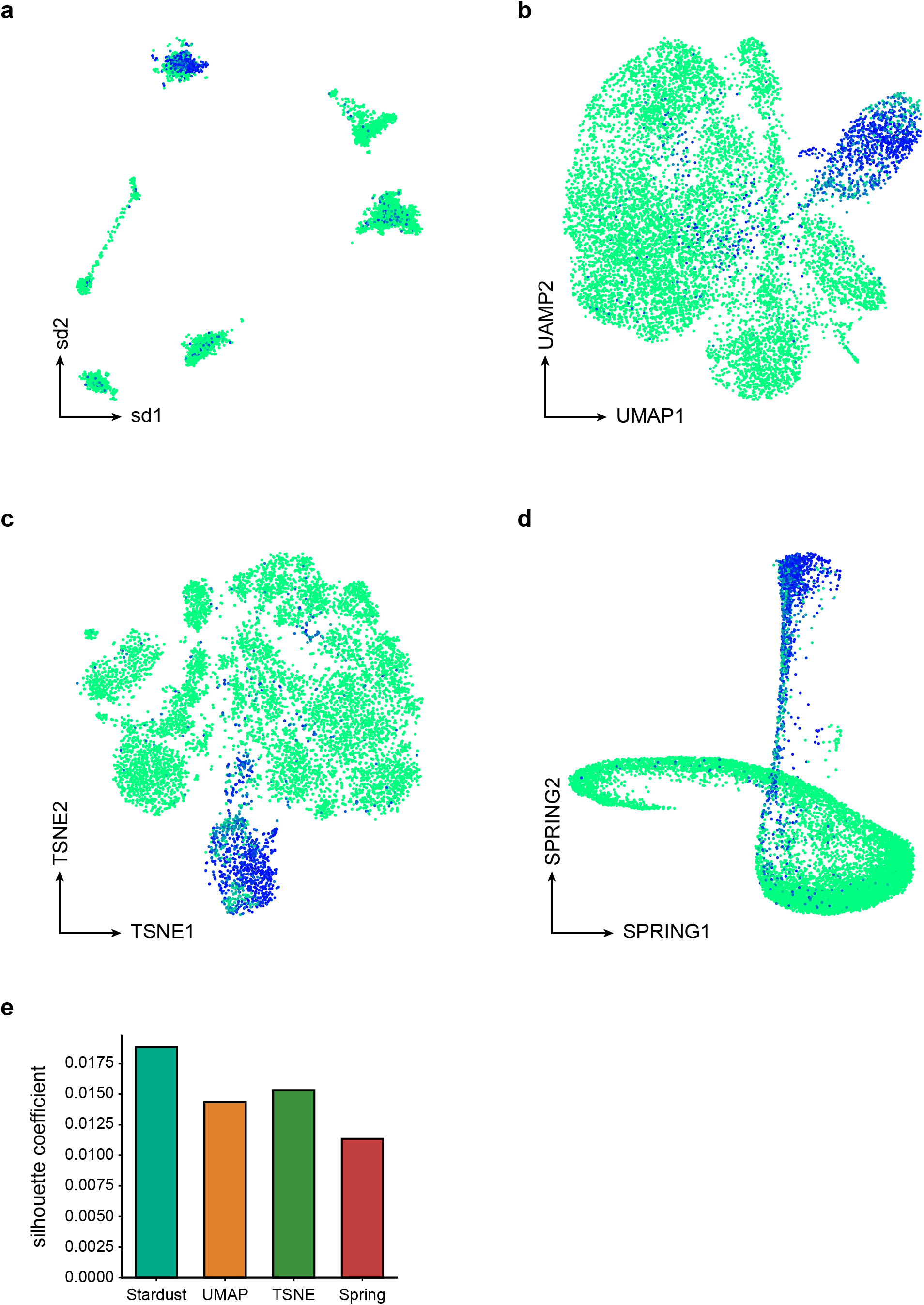
Segregation of H1N1 infected cells and bystanders. (a-d) Visualization of infected and uninfected (including bystanders and mock) cells using Stardust, t-SNE, UMAP, and SPRING. (e) Silhouette scores for the infected cells, computed based on Stardust, t-SNE, UMAP, and SPRING provided coordinates.

We quantitatively estimated these results using the Silhouette score, where Stardust scored the maximum, followed by t-SNE, UMAP, and SPRING (**Figure 2e**). Taken together, the aforementioned comparative analysis clearly advocates the outperformance of Stardust on challenging datasets harboring mixed transcriptomic features from host cells and infecting pathogens.

### Cell-type zonation in Drosophila embryo

In eukaryotes, cells often expand within a tissue niche, giving rise to cell-type zonation and patterned gene expression activities. As such, an accurate unsupervised clustering of single cell RNA sequencing profiles is expected to identify cell-groups that map with precise anatomical locations. To validate this, we considered *Drosophila* embryos, with simplistic and well-characterized morphometric features. We built on a previous work by Nitzan and colleagues [24] that successfully mapped ∼1300 single cell transcriptomes to the BDTNP virtual *Drosophila* embryos [25], using the gene expression cartography technique, while accurately preserving the spatial expression patterns of the well-known cell-lineage markers including the gap genes and the pair-rule genes. Stardust inferred 10 clusters from the scRNA-seq dataset, whereas Louvain could identify only 7 clusters (**Figure 3a, b, Supplementary Figure S7**). We visualized the clusters using Stouffer’s scores spanning cluster-specific marker genes, as well as its manually calibrated (by introducing a suitable threshold) binarized form (**Supplementary Figure S8**). Interestingly, both Louvain and Stardust exhibited a reasonable degree of zonation of the inferred cell-types and their canonical markers. However, quite evidently, the overall segregation of the spatially proximal cell-types was better in the case of Stardust. Similar results were obtained from the intercluster DE analysis (**Supplementary Figure S9**), where Stardust clustering led to the identification of marker genes, restricted to one or more clusters (**Supplementary Table 2**). As a proof-of-principle, we thoroughly investigated the anterior head region of the *Drosophila* embryo since Stardust reported a larger number of putative cell types in this region. Careful interrogation of these clusters revealed neat bifurcation of the anterior endoderm anlage and the head mesoderm by Stardust, which Louvain failed to resolve (Stardust cluster 4, and 8 vs. Louvain cluster 6) (**Figure 3c-e**). To further confirm the identity of these regions, we overlaid the publicly available in situ images available through the FlyBase webserver [26], along with the loss-of-function information of the identified markers. For example, *ImpL3* a gene, which encodes for L-lactate dehydrogenase, shows restricted expression in the anterior endoderm anlage at the embryonic stage 6 of the developing *Drosophila* larvae [27,28]. Similarly, Stardust identified *gcm*, a zinc finger transcription factor as a marker for cluster 8. This is in line with the available in situ hybridization images of *gcm* (**Supplementary Table 2**), which shows its restriction expression in the head mesoderm anlage at embryo stage 6. Of note, *gcm* is a reported marker for the head mesoderm [29], and its targets are known to regulate components of the Notch, Hedgehog (Hh), Wingless (Wg)/Wnt, Fibroblast Growth Factor (FGF) Receptor (FGFR), and JAK/STAT signaling pathways [30]. Next, as an orthogonal validation, we used probabilistic mapping scores of the top differentially expressed genes to predict the bin coordinates on the virtual embryo. Stardust produced lower values of Root Mean Squared Errors (RMSE) as compared to Louvain (**Supplementary Figure S10**). Next, we asked if Stardust inferred cell types are consistent when transcript abundance is measured using different expression assays. To this end, we compared scRNA-seq based cell clusters with the clusters of the positional bins (based on in situ expression quantification [25]). Stardust produced a highly structured similarity matrix as compared to Louvain (**Supplementary Figure S11**).

**Figure 3.**
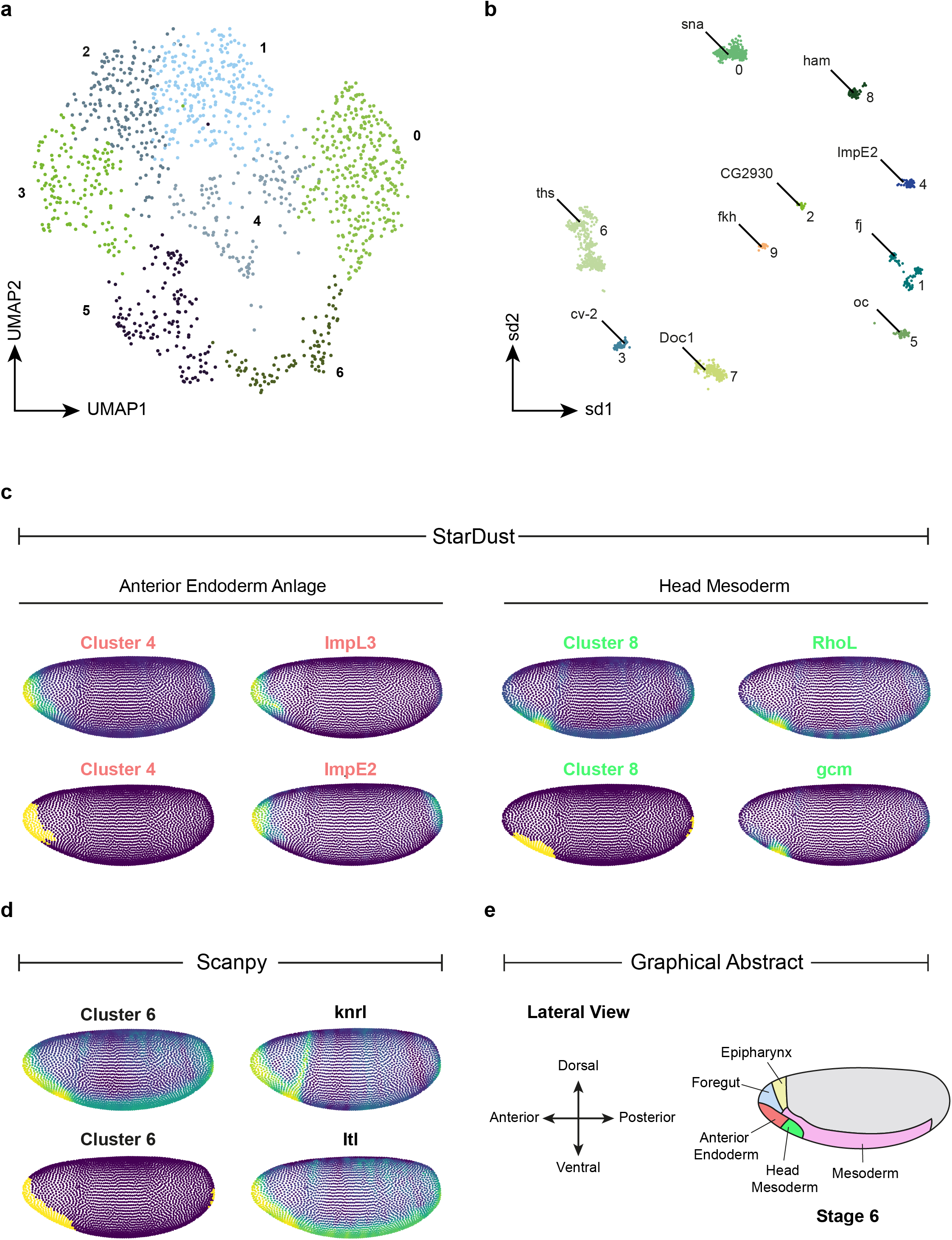
Precise delineation of cell-types in *Drosophila* early embryos. (a) UMAP based visualization of cell-type clusters obtained using scanpy. (b) Visualization of cell-type clusters and marker genes using Stardust. (c) Location of Stardust clusters 4, and 8, and the associated marker genes on *Drosophila* virtual embryo (locations are marked based on Stouffer’s score gradient, as well as its binarised form as discussed in the Online Methods section). (d) Equivalent plots for Louvain cluster 6. (e) Schematic diagram representing the lateral view of the stage 6 *Drosophila* larva with color-coded key anatomical regions.

Stardust employs an iterative cell-gene co-embedding algorithm that mirrors the spirit of an expectation-maximization process. It consumes significantly more time as compared to the existing best practice methods. For instance, in the case of ∼68K *PBMC* data Stardust took 1 hour 48 minutes to complete the analysis, whereas Louvain took only 3.26 minutes. On the positive side, Stardust offers radically cleaner embedding of single cells, as compared to the state of the art, and allows to visualize marker genes alongside single cells. Further, its ability to distinguish functionally distinct cell populations with overlapping gene expression patterns makes it a promising alternative to the existing pipelines for scRNA-seq data analysis.

## MATERIALS AND METHODS

### Description of the datasets

We worked with seven different single cell expression datasets to evaluate the Stardust for various analyses. The first dataset comprises ∼68,000 peripheral blood mononuclear cell (PBMC) transcriptomes from a healthy donor. The authors annotated the cells by correlating their expression profiles with 11 common immune cell subtypes purified using Fluorescence-Activated Cell Sorting (FACS). This dataset served as a gold standard for comparative analysis of the clustering techniques. The second data set consists of host and viral transcriptomes of about 9000 individual cells, infected with single influenza A virus (IAV) [23]. This dataset is used for comparative analysis of the visualization techniques. The third dataset consists of ∼20k single cell transcriptomes sampled from the arcuate-median eminence complex (Arc-ME) region of mouse brain [22]. The fourth dataset consists of 4645 single cell expression profiles sampled from melanomas of 19 patients [31]. Based on the available annotations we discarded the clusters with cluster-specific cell populations less than 50. Fifth data single cell RNA sequencing dataset of mouse embryonic stem cells consists of 301 single cells from 11 populations [32]. Sixth dataset consists of 704 single cell transcriptomes of pluripotent stem cell-derived cerebral organoids from chimpanzees [33]. The seventh dataset is a scRNA-Seq data of the human liver comprising about ∼7000 cells [21]. We used spatially resolved single cell transcriptomes of *Drosophila* embryo [34] The scRNA-Seq data reported in this study consists of about 1300 high-quality single cells having >12,500 unique transcripts and expressing more than five genes of the Berkeley *Drosophila* Transcription Network Project (BDTNP) reference atlas [25] expressed over ∼8000 genes. Karaiskos and colleagues putatively resolved the spatial location of the profiled single cells by quantifying expression similarity across single cells and ∼3000 positional bins. Notably, the BDTNP reference atlas quantifies the relative mRNA levels of 84 genes in a virtual stage 5 *D. melanogaster* embryo [25].

### Preprocessing of the scRNA-Seq data

Expression matrices are loaded as AnnData objects, as implemented in the scanpy pipeline [35]. Cells expressing less than 30% of the average number of expressed genes across cells are filtered. Further, we retained those genes which are detected in at least 3 cells. The filtered dataset is subjected to Unique Molecular Identifier (UMI) count normalization where the counts per cell are normalized to the median of the total counts of all the cells before normalization. The normalized expression matrix is further log-transformed after adding 1 as pseudo-count. Percentage of the highly variable genes based on the size of the dataset is selected from the preprocessed expression matrix using the scanpy function i.e. *sc*.*pp*.*higly_variable_genes()* with the parameter *n_top_genes* set to 500. The function internally uses Cell Ranger for computing normalized dispersion.

### Stardust outline

Stardust works by constructing a network of single cells and genes such that expression-wise similar cells are connected with each other, and genes are connected to cells where those are enriched the most. This gives rise to a cell-gene multi-entity network, which is subjected to a graph layouting algorithm, which provides 2D coordinates of both cells and the genes in a manner that cells that share their lineage and their marker genes maintain proximity. At this stage, cells and genes are clustered based on the coordinates associated with their low-dimensional embeddings. Differentially Expressed (DE) genes are obtained by contrasting the cell-clusters. The DE genes replace the highly variable genes used for the construction of the initial network. This step is followed by layouting for further refinement of the 2D embedding. Typically, 3 iterations of this refinement process offer an optimal representation of the cells. The subsequent sections outline the details associated with the main components of Stardust.

### Structure Preserving Sub-sampling

Random subsampling might result in the loss of rare sub-populations. To circumvent this, Stardust employs Structure Preserving Sampling (SPS) [5] on the preprocessed datasets. SPS involves rapid construction of the nearest neighbor network of single cells, followed by graph clustering. To ascribe a higher retention rate to rare cells, cell sub-sampling is done based on the following decay function, which determines the proportion of cells to be sampled per cluster.

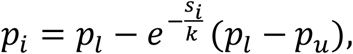

where *p*_*i*_ represents the proportion of samples for *i*^*th*^ cluster, *p*_*l*_ and *p*_*u*_ represents the lower bound and upper bound proportion respectively, *s*_*i*_ is the size of the cluster and *k* is the scaling factor.

### Cell-gene Nearest Neighbor Network (CGNNN)

The sampled cells after SPS and the qualified genes based on the gene selection technique (discussed earlier) conjointly construct the CGNNN. A CGNNN consists of two legal types of edges - cell-cell and cell-gene. For cell-cell connections, Principal Component Analysis (PCA) is performed on the preprocessed, normalized and log transformed expression matrice. The default number of top components is set to 500. The concept of approximate nearest neighbor is used for establishing cell-cell connections by employing Locality Sensitive Hashing (LSH). Each cell is connected to its *k* nearest cells based on Annoy (Approximate Nearest Neighbors Oh Yeah) a C++ library with Python bindings, on the basis of top principal components. The construction of the Nearest Neighbor Network (NNN) takes *O*(*nlog n*) where *n* denotes the number of single cell expression profiles. The usage of LSH results in reduction of the computation time of NNN. Similarly, each gene is connected to top *n* cells in terms of normalized expression estimates. The default values of *k* and *n* are set to 20 and 10 respectively. The count of qualified genes per pass for construction of CGNNN is described in detail in the section titled “OpenOrd based iterative graph layouting”. As the number of qualified genes is lesser in the 4th pass as compared to other passes, so the number of cell-gene connections is slightly different. In the 4^th^ pass, each cluster-specific gene identified in the 3^rd^ pass at rank *i* in the cluster is connected to *p* cells of the same cluster s.t. *p* = 5 − ⌈*i*/10⌉. The intent here is to attach cluster-specific genes of higher quality to more cells within the cluster.

### Post hoc cluster assignment

In every pass, density-based clustering (DBSCAN) is performed on coordinates corresponding to 2D embeddings of the subsampled transcriptomes. Cells leftover during the subsampling phase are assigned post-hoc, to the predefined clusters. For each of the leftover transcriptomes, *k* nearest neighbours are queried from the subsamples selected for the initial clustering. A leftover transcriptome is assigned to the cluster which is found maximally represented among the neighbors. The default value of the parameter *k* is set to 20, the same as that of the cell-cell connections parameter described in the CGNNN section. The 2D coordinates of the leftover cells are determined by averaging out the coordinates of the nearest neighbor transcriptomes residing inside the assigned cluster. Notably, the post-hoc cluster assignment is performed at the end of each pass, which helps in determining the coordinates and clusters of origin for the leftover cells. The only exception is the 4^th^ pass where cluster assignment is not necessary.

### OpenOrd based iterative graph layouting

Stardust pipeline involves iterative embedding and differential gene finding with a goal of converging into high-quality cell-type clusters. In the 1^st^ pass, 500 genes with the highest absolute loadings across top 500 PCs are selected to construct the CGNNN. In the 2^nd^ pass and 3^rd^ pass, using scanpy’s *scanpy*.*tl*.*rank_genes_groups()* with *t-test_overestim_var* as the underlying method, 500 differentially expressed genes with adjusted *p-value <* 0.5 and *log2foldchange* > 1.2 are selected for CGNNN. In 4^th^ pass, the top 20 cluster-specific differentially upregulated genes are considered for reconstruction of the CGNNNs. The multi-entity, heterogeneous network created in every pass is subjected to OpenOrd network layouting.O penOrd algorithm uses simulated annealing for optimization of the below expression.

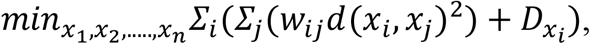

where *x*_*i*_ represents the position of the node, 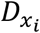 denotes the density of the edges near *x*_*i*_. *w*_*ij*_ denotes the edge weight connecting node *i* and *j*. Finally, *d*(*x*_*i*_, *x*_*j*_) denotes the distance between the two nodes. Notable, OpenOrd uses an edge-cutting heuristic that allows the user to control the amount of white space. Stardust considers unit weights for all the connections (edges). The low-dimensional embedding of single cell transcriptomes obtained in the 3^rd^ pass are used as input dimensions for clustering. Final coordinates of the cells and genes are determined through one last round of co-embedding in the 4^th^ pass (see the subsection titled “Final coordinates for differential genes”).

### Final coordinates for differential genes

Cluster-specific top differentially upregulated genes obtained through DE analysis in the 3^rd^ pass are subjected to CGNNN formation for one final time. The detail for the NNN formation in the 4^th^ pass is described in the section titled “Cell-gene Nearest Neighbor Network (CGNNN)”. Execution of the OpenOrd graph layouting algorithm on the multi-entity network yields a new set of coordinates for the cells and the genes. The cell coordinates may vary from the 3^rd^ pass coordinates which we claim to be the final coordinates of the transcriptomes. 4^th^ pass helps in mapping newly obtained cluster-specific upregulated genes back into the 2D space generated in the 3^rd^ pass. This is achieved by training cluster-specific regression models wherein each dimension obtained from the 3^rd^ pass is considered as the dependent variable, whereas the two dimensions obtained from the 4^th^ pass are considered as the independent variables. For genes specific to each cluster obtained in the 3^rd^ pass, the corresponding coordinate values are generated using these models on the basis of the final embedding in the 4^th^ pass. Non-linear K Nearest Neighbor (KNN) regression technique is used for this purpose. The default value of K is set to 5. To simplify, both 2D coordinates and clusters are obtained from the 3^rd^ pass. 4^th^ i.e. the final pass is used only for determining the putative location of the cluster-specific differentially upregulated genes in the 2D space generated in the 3^rd^ pass.

### Projection of DE genes onto UMAP embedding

As an alternative to the OpenOrd layout, Stardust also allows cell-gene co-embedding on the UMAP projections. This is achieved simply by projecting the cluster-specific genes obtained from the 3^rd^ pass onto the UMAP laid 2D space. The mapping strategy is similar to the 4^th^ to 3^rd^ pass mapping technique described in the previous section. In this case, the UMAP dimensions individually are considered as the decision variables for regression.

### Measures of clustering performance

We used both unannotated and annotated scRNA-seq datasets for various experiments. For assessing the signal-to-noise ratio associated with the representation of clusters on 2D maps, we made use of the Silhouette index.

### Analysis of spatial transcriptomics data of *Drosophila* embryo

As discussed earlier, we used independent single cell expression data [36] and in situ expression estimates [25] of *Drosophila* larvae to assess the performance of Stardust and Louvain. The in situ data was available as ∼3000 positional bins expressing 84 marker genes. We performed separate analysis of in situ and the single cell expression data using both Louvain and Stardust. To identify the correspondence among clusters obtained from the transcriptomics data and the in situ hybridization data, we tracked the cluster-to-cluster correspondence by computing Pearson’s correlation between cluster-wise average expression vectors of the common 84 genes, present in both datasets.

### Spatial zonation of cell-type clusters in *Drosophila* embryo

To track the spatial localization of the clusters generated using Stardust and Louvain, we made use of the gene expression cartography technique, implemented in the novoSpaRc [24] framework. NovoSpaRc probabilistically mapped the ∼8,000 genes in the scRNA-seq dataset across positional bins (coordinates of the bins were supplied as input) with the underlying assumption that relatively closer cells by position tend to share similar gene expression patterns and vice versa. These probabilities mapping scores can be treated as a surrogate of gene expression estimates across the positional bins. To pinpoint the putative spatial localization of a single cell cluster, we computed Stouffer’s Z score by combining the probabilistic mapping scores associated with the top 20 cluster-specific differentially upregulated genes. This allowed us to obtain cluster-specific continuous scores across positional bins of the virtual *Drosophila* embryo.

### Prediction of spatial coordinates using differentially expressed genes as features

The ∼8K X ∼3000 matrix returned by novoSpaRc comprises probabilistic mapping scores of ∼8K genes across ∼3000 positional bins. For Stardust, and Louvain, we selected a total of ∼100 DE genes, while considering an equal number of marker genes per cluster. We constructed 50 bootstrapped samples of training and validation sets by sampling from the positional bins. *DecisionTreeRegressor* module of *sklearn* was used for the prediction tasks while treating the marker genes as independent and each of the coordinates as a dependent variable. The performance of the various regression tasks was tracked using Root Mean Square Error (RMSE).

## CONCLUSION

The declining cost of sequencing and wide availability of single cell omics platforms have created an information singularity for studying cellular processes under normal and diseased conditions. High throughput capture and profiling of single cells enable characterizing tens of cell types and states as part of a single experiment. Current clustering and differentially expressed gene detection are treated as independent processes, thereby generating independent 2D scatters and heatmaps respectively. Cross-referencing between these two types of visualizations is often challenging. Moreover, cell-type clusters are often detected with diffused identities. Stardust addresses these by iteratively clustering cells and marker genes as part of multi-entity network topology. Stardust successfully groups cells and genes together in a visually coherent manner. Stardust’s iterative clustering yields cell type clusters with high signal-to-noise ratios. With the increasing throughput of single cell experiments, we predict Stardust would emerge as a non-redundant alternative, especially due to its compatibility with common single cell visualization techniques such as UMAP and tSNE.

## SOFTWARE AVAILABILITY

Stardust is freely available at https://github.com/Swagatam123/Stardust_package as a python package.

## AUTHOR CONTRIBUTION

DS conceived the study and supervised the method development, with inputs from GA and TC. GA supervised the designing of experiments with inputs from DS. SC implemented the software pipeline and conducted all experiments with help from NB, KG, D.Sinha, and AM. AM performed post-processing of the scientific illustrations. The manuscript was written by DS, SC, NB, CN, and GA. All the authors read and approved the manuscript.

## ACKNOWLEDGEMENT

The Sengupta lab is funded by the INSPIRE faculty grant from the Department of Science & Technology (DST), India. The Ahuja lab is supported by the Ramalingaswami Re-entry Fellowship by the Department of Biotechnology (DBT), Ministry of Science & Technology, Govt. of India.

