## Supplementary Information for "Simultaneous visualization of cells and marker genes from scRNA-seq studies"

**Supplementary Figure S1.** Steps involved in Stardust based co-embedding of cells and genes.

**Supplementary Figure S2.** (a) Distribution of annotated melanoma cells across clusters obtained using Stardust. (b) Equivalent plots for Louvain based clusters.

**Supplementary Figure S3.** (a) UMAP based visualization of the *PBMC* data. The red triangles on the 2D map show the coordinates of the cluster specific upregulated genes. The top differentially upregulated marker gene is shown per cluster. (b) Stardust based co-embedding of cells and genes from the *Melanoma* dataset. Coordinates of the marker genes are not plotted due to smaller cluster sizes and higher cellular density. (c) Alluvial plot depicting the correspondence between unsupervised clusters obtained using both the methods, and the cell annotations reported by Zheng and colleagues(28).

**Supplementary Figure S4.** (a) UMAP based visualization of the liver data(18), with marker gene locations inferred by Stardust. The red triangles on the 2D map show the coordinates of the cluster specific upregulated genes. (b) Stardust based co-embedding of cells and genes from the *Melanoma* dataset. Coordinates of the marker genes are not plotted due to smaller cluster sizes and higher cellular density.

**Supplementary Figure S5.** (a) UMAP based visualization of the mouse brain data(19), with marker gene locations inferred by Stardust. The red triangles on the 2D map show the coordinates of the cluster specific upregulated genes. (b) Stardust based co-embedding of cells and genes from the Melanoma dataset. Coordinates of the marker genes are not plotted due to smaller cluster sizes and higher cellular density. (c) Alluvial plot depicting the correspondence between unsupervised clusters obtained using LouvainStardust and authors' annotation of the Seurat(35) clusters.

**Supplementary Figure S6.** Cluster specific and average Silhouette scores computed for each dataset using both Louvain and Stardust based on cell coordinates on the 2D maps.

**Supplementary Figure S7.** (a) Spatial zonation of Stardust-derived clusters is shown on the virtual *Drosophila* embryos (lateral view). Color coding is done based on Stouffer's scores entailing cluster-specific marker genes. Bright yellow indicates a higher overall expression of the transcripts, whereas dark blue indicates regions with limited or no enrichment. (b) Similar plots for Louvain derived clusters.

**Supplementary Figure S8.** (a) Spatial zonation of Stardust-derived clusters is shown on the virtual *Drosophila* embryos (lateral view). Color coding is done based on binarized Stouffer's scores entailing cluster-specific marker genes. The bright yellow patches indicate spatial zonation of the clusters on a virtual *Drosophila* embryo. (b) Similar plots for Louvain derived clusters.

**Supplementary Figure S9.** (a) Heatmap of the cluster specific top differentially expressed genes identified from the scRNA-seq data of *Drosophila*, considering the clusters inferred by Stardust. (b) Similar heatmap for Louvain derived cluster specific differentially expressed genes.

**Supplementary Figure S10.** (a) RMSE score comparison between Stardust and Louvain across multiple bootstrap runs on predicting x-coordinate on the virtual *Drosophila* embryo. (b) RMSE score comparison between Stardust and Louvain across multiple bootstrap runs on predicting y-coordinate on the virtual *Drosophila* embryo. (c) RMSE score comparison between Stardust and Louvain across multiple bootstrap runs on predicting z-coordinate on the virtual *Drosophila* embryo.

**SupplementaryFigure S11.** (a) Heatmap of the Pearson's correlation between clusters obtained by Stardust from scRNA-seq data and in situ data of *Drosophila* embryos. Significant correspondence is visible between the clusters. (b) Similar heatmap for the Louvain derived clusters. (c) For each of the Stardust derived clusters from scRNA-seq data, we plotted the distribution of Pearson's correlation with the in situ based clusters. Majority of the clusters exhibited a bimodal distribution indicating their selective similarity with a subset of clusters obtained by analyzing the in situ based expression estimates.

(d) Similar density plots for Louvain derived clusters. In this case, clear bimodal distribution is observed in a rather small number of cases.

**Supplementary Table 1.** List of cluster specific differentially upregulated genes retrieved using Stardust and Louvain on various datasets.

**Supplementary Table 2.** Links to in situ images corresponding to top marker genes for the clusters obtained using Stardust and scanpy by analyzing the *Drosophila* scRNA-seq dataset.

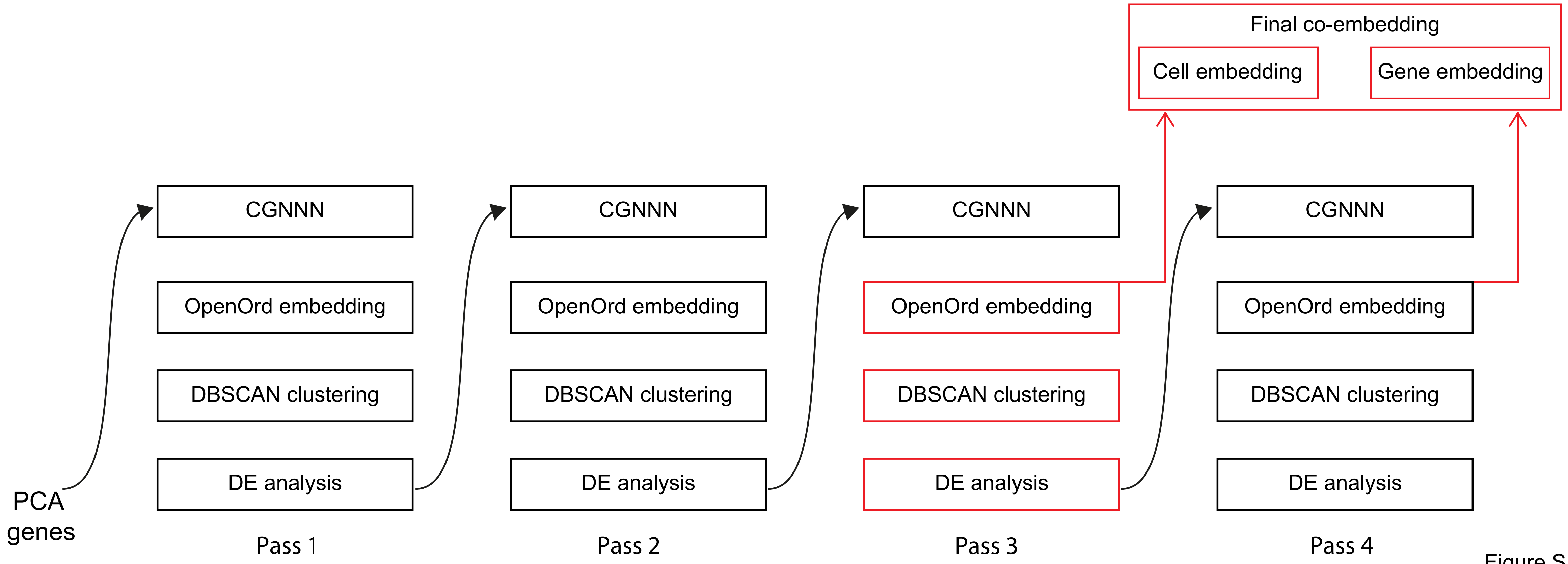
Final co-embedding
Cell embedding
Gene embedding
CGNNN
OpenOrd embedding
DBSCAN clustering
DE analysis
CGNNN
OpenOrd embedding
DBSCAN clustering
DE analysis
CGNNN
OpenOrd embedding
DBSCAN clustering
DE analysis
CGNNN
OpenOrd embedding
DBSCAN clustering
DE analysis
PCA genes
Pass 1
Pass 2
Pass 3
Pass 4

Figure S1

**a**

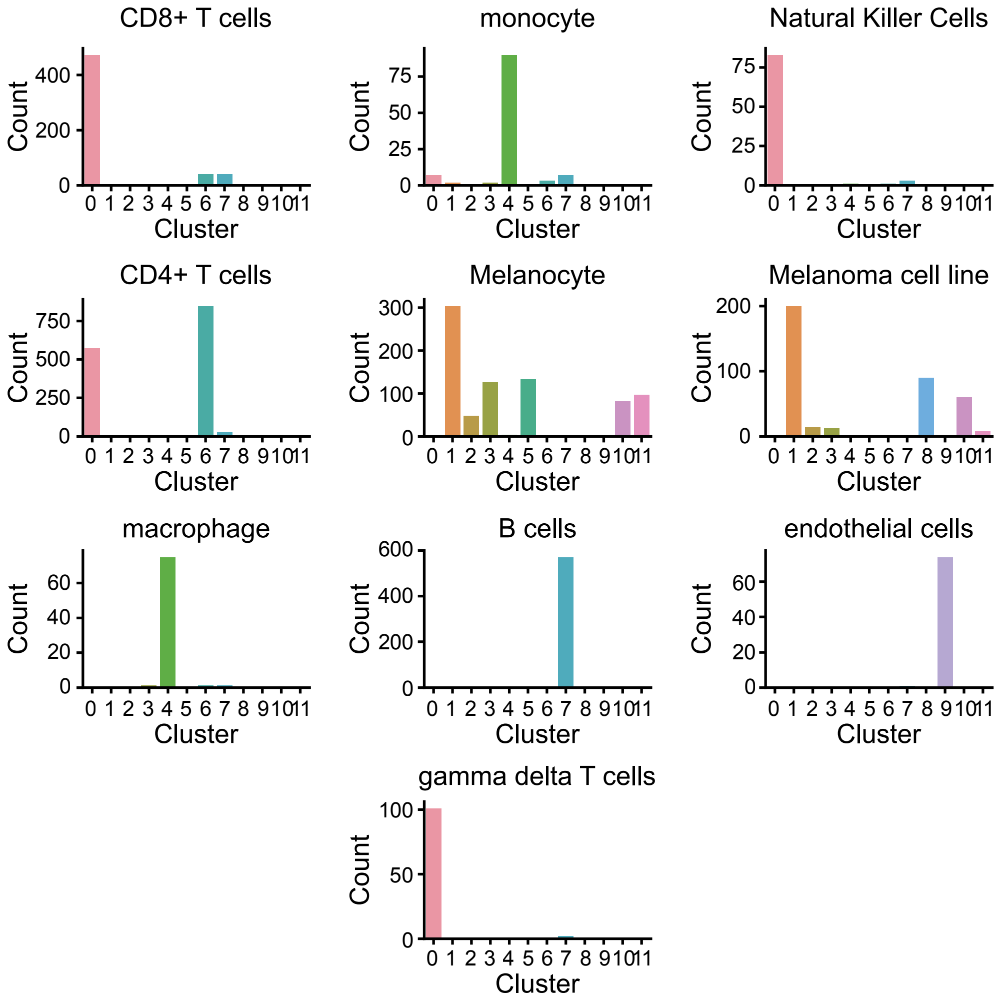


Correspondence between annotations and Stardust clusters

**b**

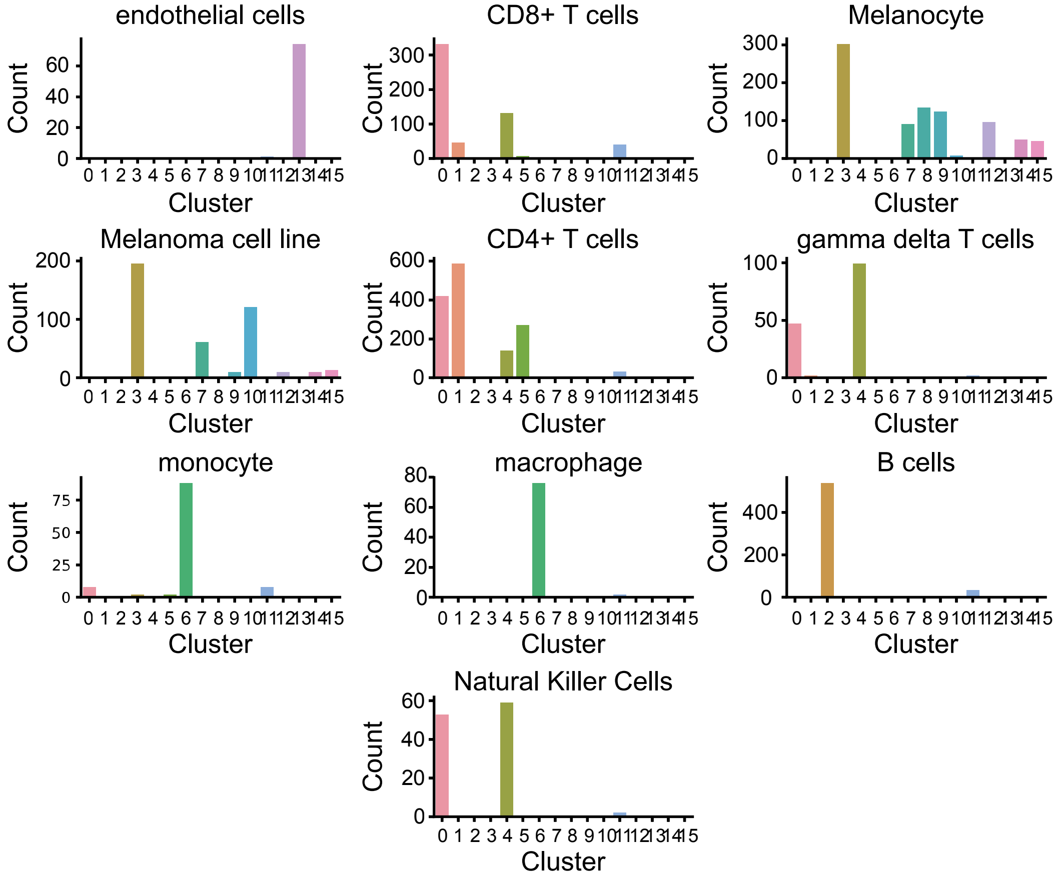


Correspondence between annotations and Louvian clusters

Figure S2

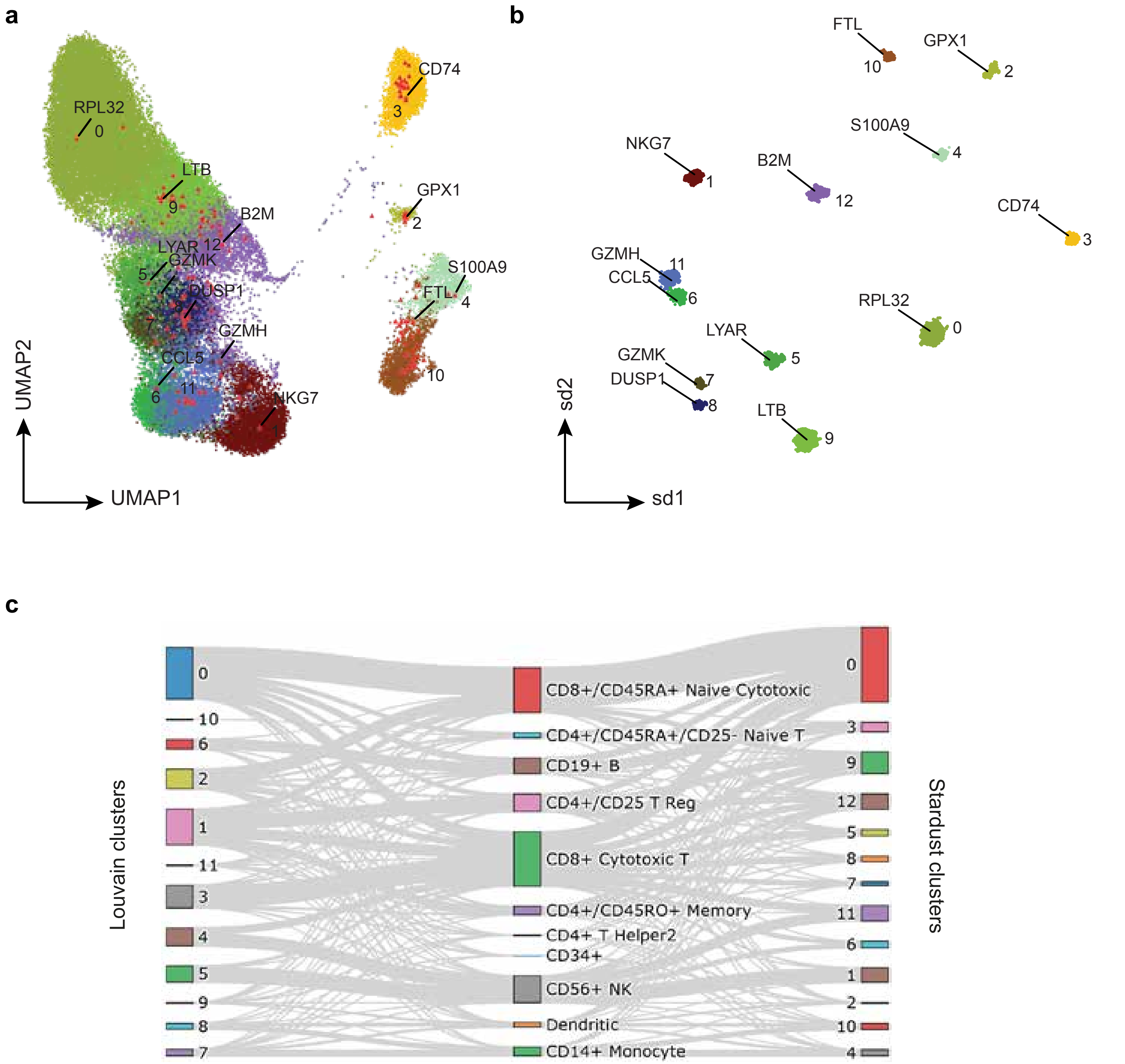
a
RPL32
LTB
B2M
LYAR
GZMK
DUSP1
GZMH
CCL5
NKG7
CD74
GPX1
S100A9
FTL
UMAP2
UMAP1
b
FTL
GPX1
S100A9
NKG7
B2M
CD74
GZMH
CCL5
RPL32
LYAR
GZMK
DUSP1
LTB
sd2
sd1
c
Louvain clusters
CD8+/CD45RA+ Naive Cytotoxic
CD4+/CD45RA+/CD25- Naive T
CD19+ B
CD4+/CD25 T Reg
CD8+ Cytotoxic T
CD4+/CD45RO+ Memory
CD4+ T Helper2
CD34+
CD56+ NK
Dendritic
CD14+ Monocyte
Stardust clusters

Figure S3

a

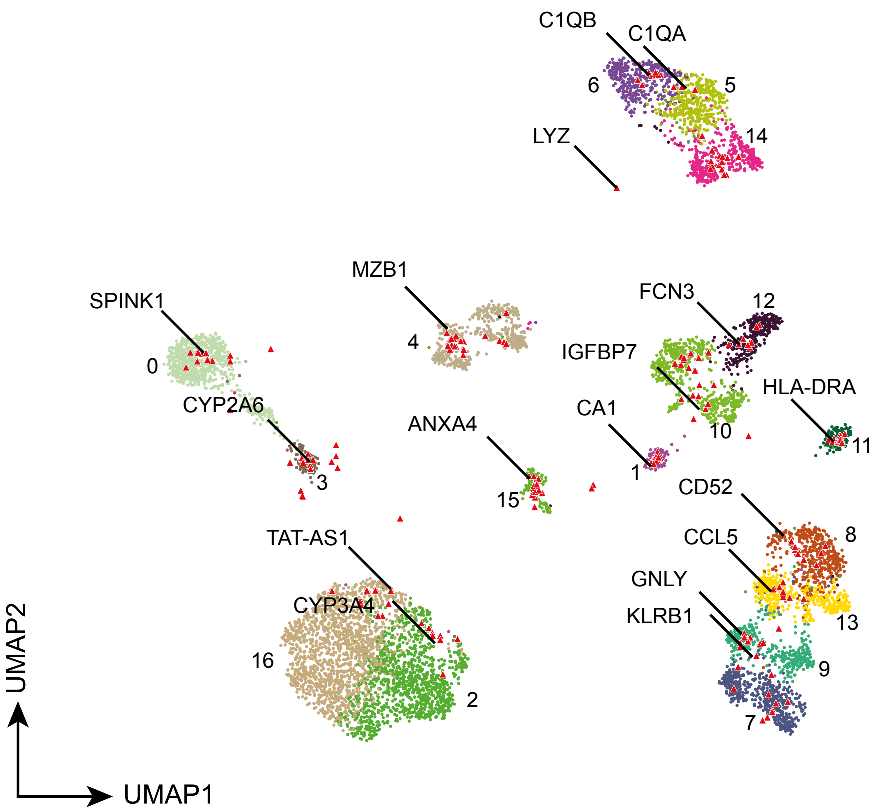
C1QB
C1QA
6
5
14
LYZ
MZB1
FCN3
12
SPINK1
4
IGFBP7
0
CYP2A6
HLA-DRA
CA1
10
ANXA4
11
1
3
15
CD52
TAT-AS1
CCL5
8
GNLY
KLRB1
CYP3A4
13
16
9
2
7
UMAP2
UMAP1

b

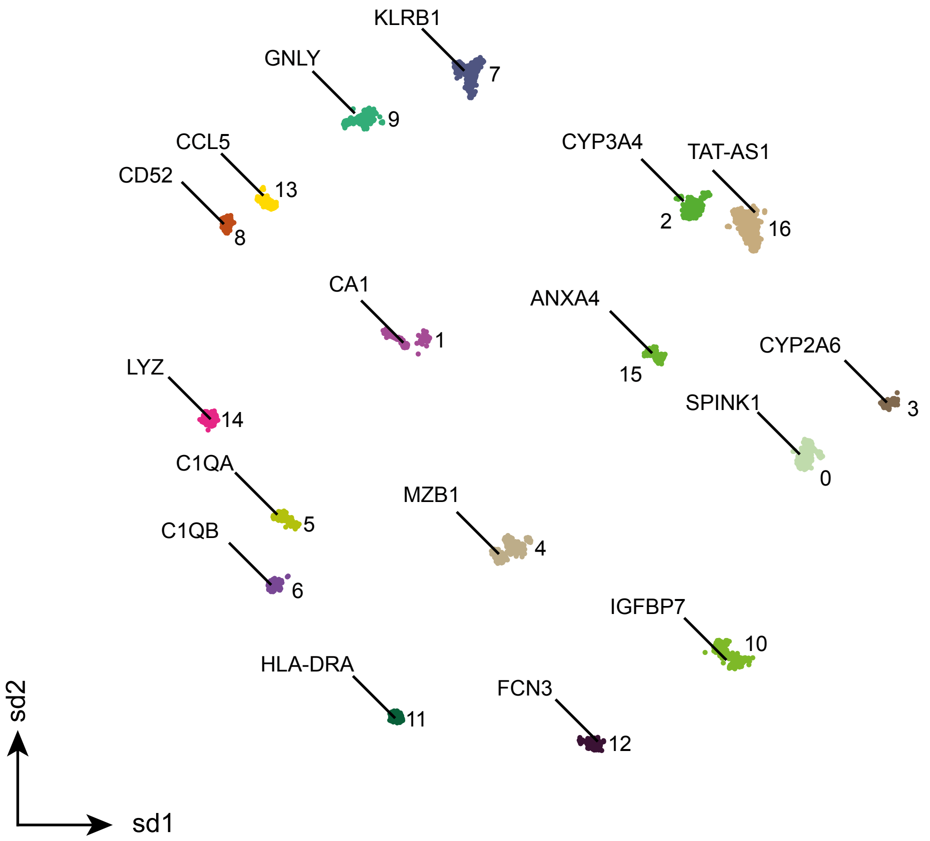
KLRB1
GNLY
7
9
CCL5
CYP3A4
TAT-AS1
CD52
13
2
16
8
CA1
ANXA4
1
CYP2A6
LYZ
15
SPINK1
3
14
0
C1QA
MZB1
5
C1QB
4
6
IGFBP7
HLA-DRA
10
FCN3
11
12
sd2
sd1

Figure S4

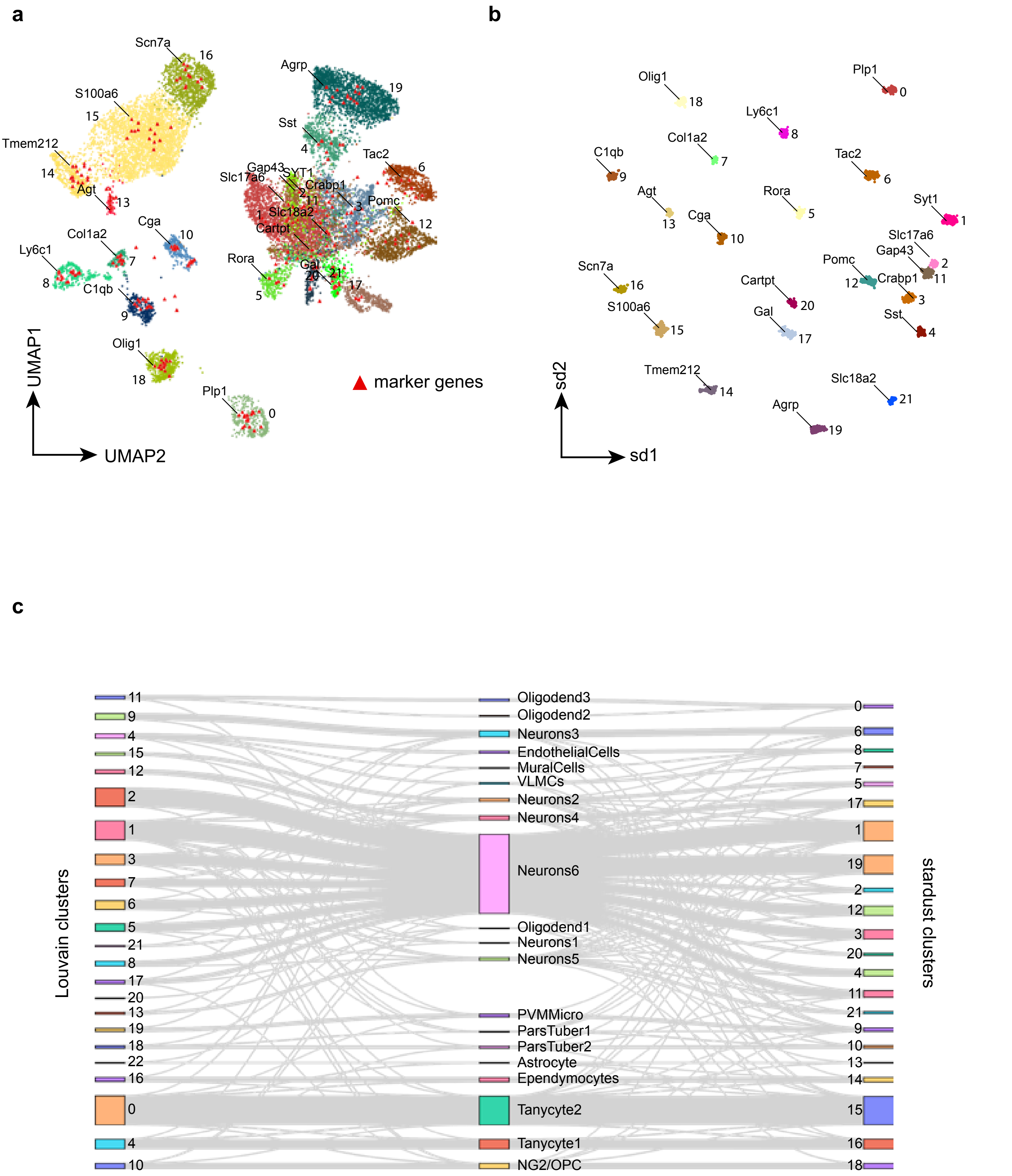
a
b
c
marker genes
UMAP1
UMAP2
sd1
sd2
Louvain clusters
stardust clusters
Oligodend3
Oligodend2
Neurons3
EndothelialCells
MuralCells
VLMCs
Neurons2
Neurons4
Neurons6
Oligodend1
Neurons1
Neurons5
PVMMicro
ParsTuber1
ParsTuber2
Astrocyte
Ependymocytes
Tanycyte2
Tanycyte1
NG2/OPC

Figure S5

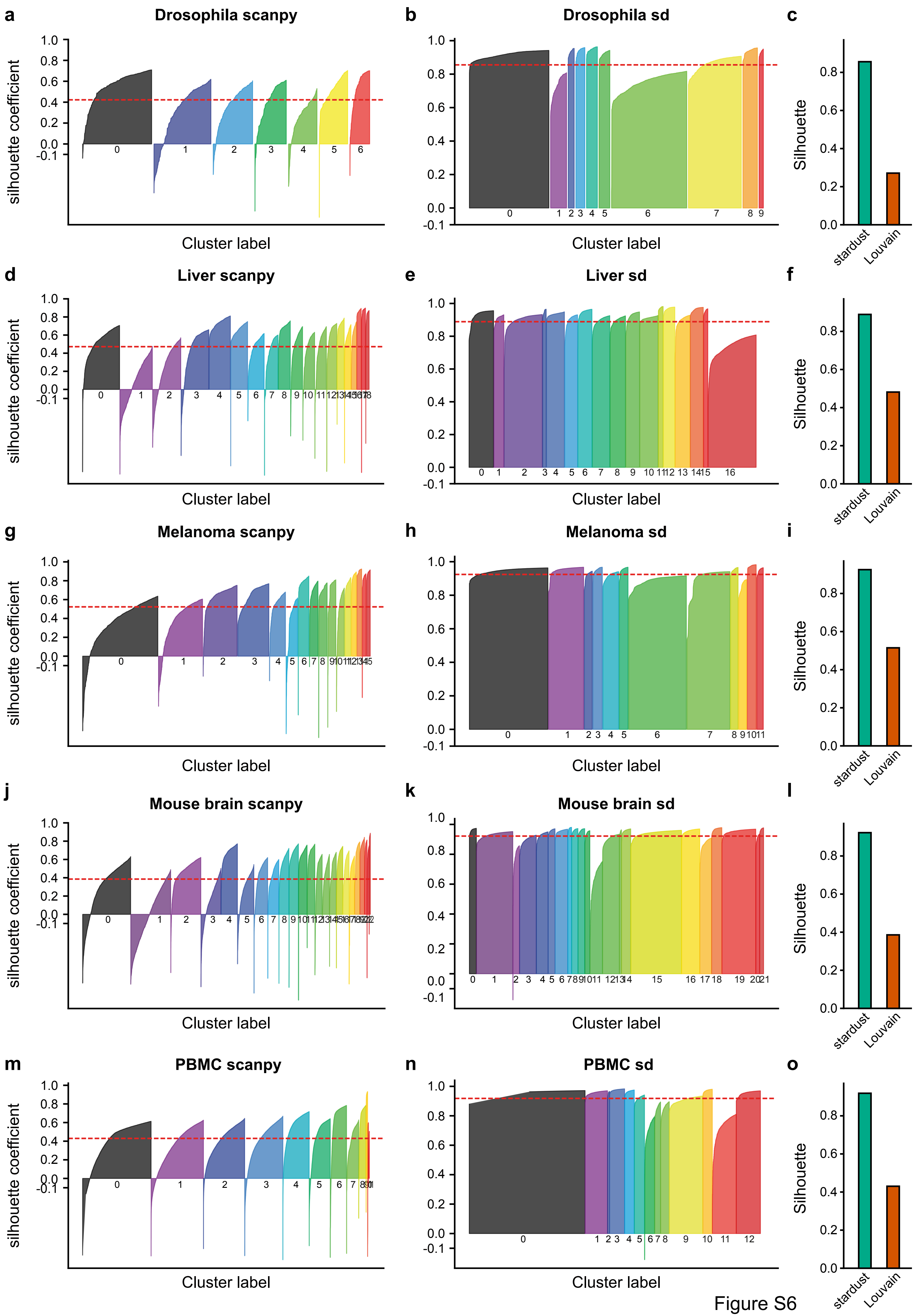

a
Drosophila scanpy
b
Drosophila sd
c
d
Liver scanpy
e
Liver sd
f
g
Melanoma scanpy
h
Melanoma sd
i
j
Mouse brain scanpy
k
Mouse brain sd
l
m
PBMC scanpy
n
PBMC sd
o
silhouette coefficient
Cluster label
Silhouette
stardust
Louvain


Figure S6

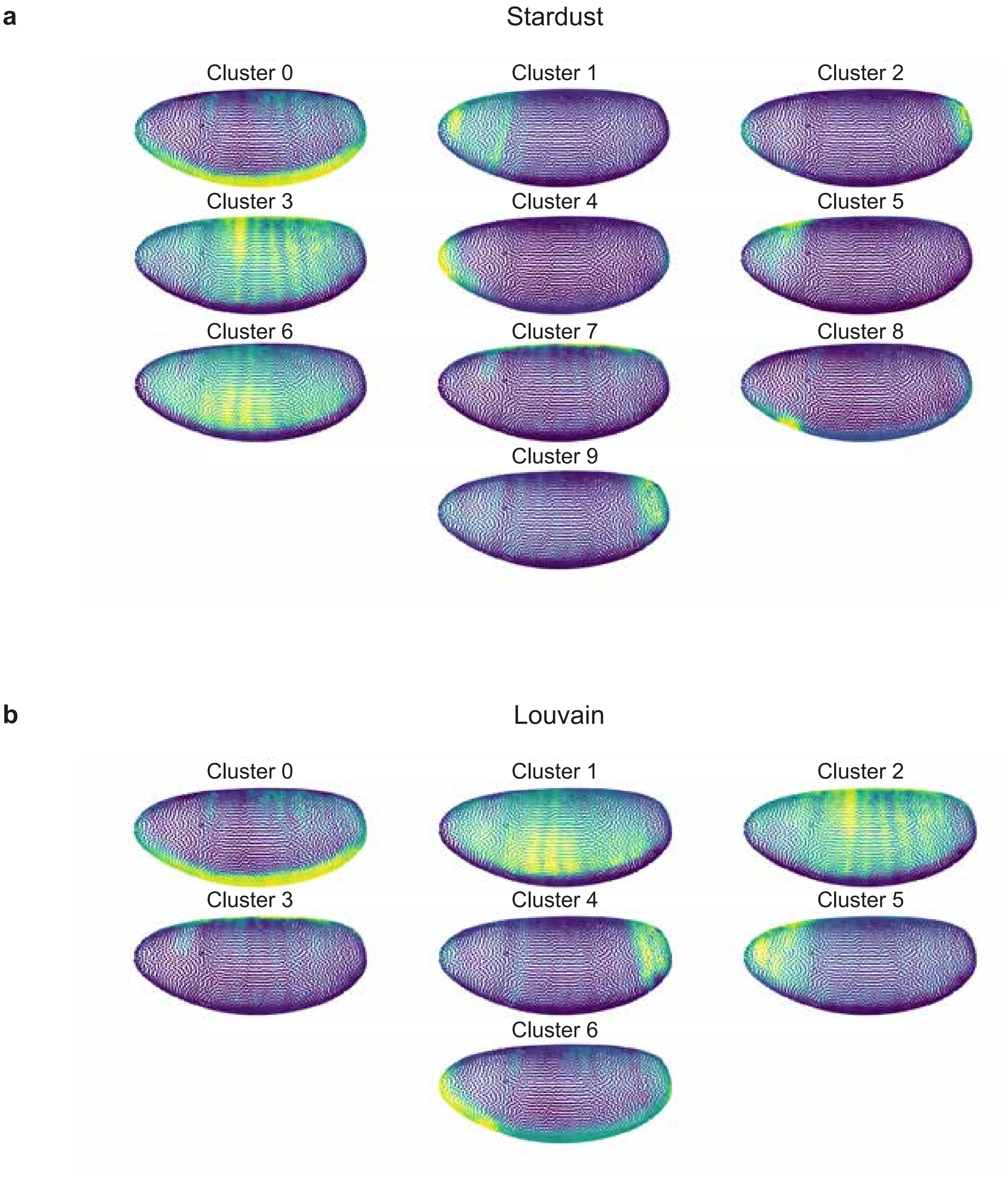
a
Stardust
Cluster 0
Cluster 1
Cluster 2
Cluster 3
Cluster 4
Cluster 5
Cluster 6
Cluster 7
Cluster 8
Cluster 9
b
Louvain
Cluster 0
Cluster 1
Cluster 2
Cluster 3
Cluster 4
Cluster 5
Cluster 6

Figure S7

**a**

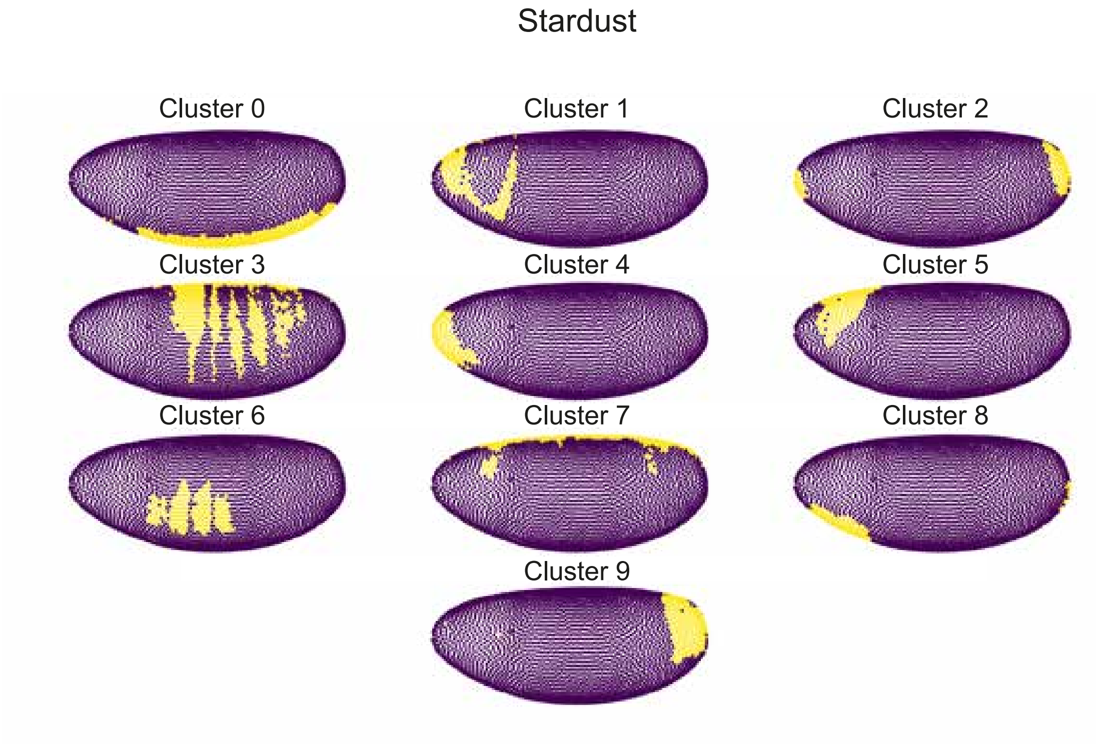
Stardust
Cluster 0
Cluster 1
Cluster 2
Cluster 3
Cluster 4
Cluster 5
Cluster 6
Cluster 7
Cluster 8
Cluster 9

**b**

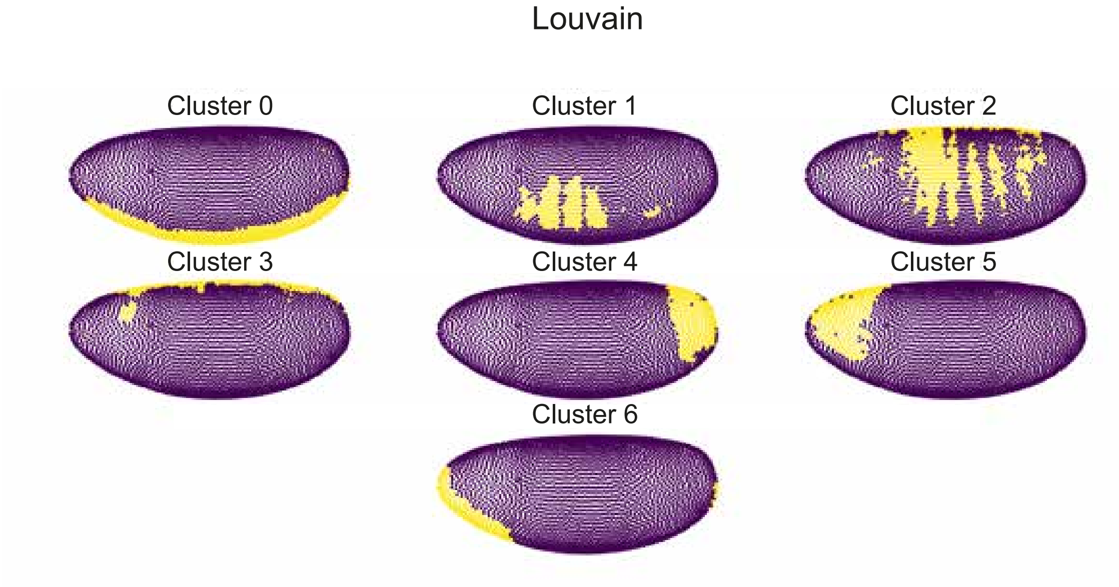
Louvain
Cluster 0
Cluster 1
Cluster 2
Cluster 3
Cluster 4
Cluster 5
Cluster 6

Figure S8

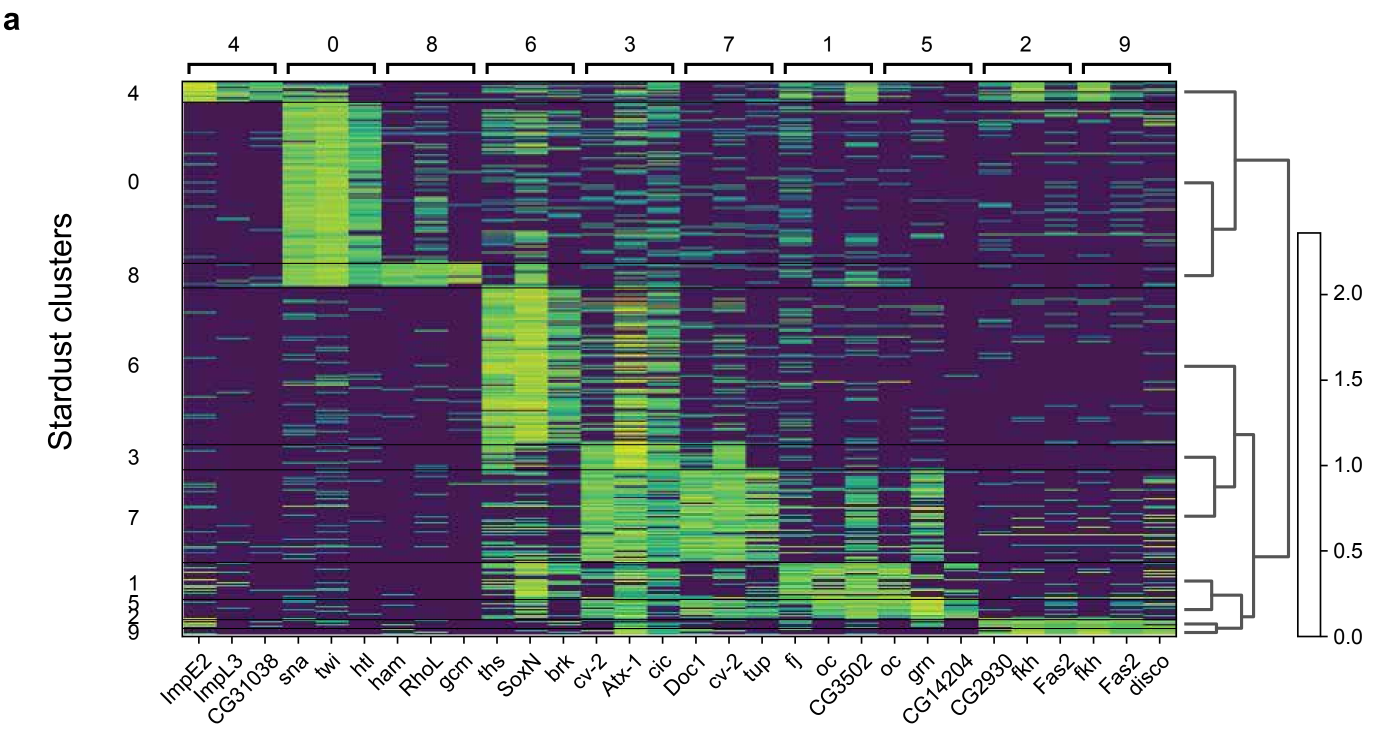

a
4
0
8
6
3
7
1
5
2
9
Stardust clusters
ImpE2
ImpL3
CG31038
sna
twi
htl
ham
RhoL
gcm
ths
SoxN
brk
cv-2
Atx-1
cic
Doc1
cv-2
tup
fj
oc
CG3502
oc
grn
CG14204
CG2930
fkh
Fas2
fkh
Fas2
disco
2.0
1.5
1.0
0.5
0.0


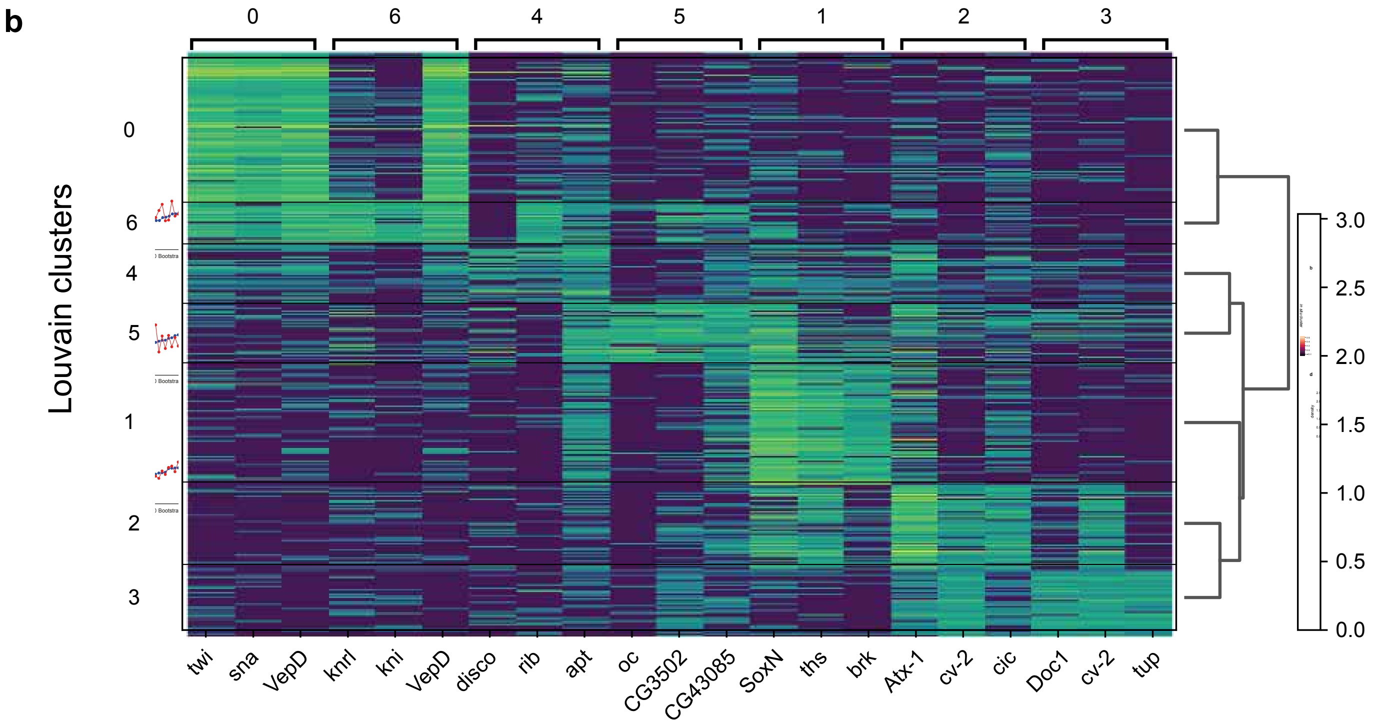

b
0
6
4
5
1
2
3
Louvain clusters
twi
sna
VepD
knrl
kni
VepD
disco
rib
apt
oc
CG3502
CG43085
SoxN
ths
brk
Atx-1
cv-2
cic
Doc1
cv-2
tup
3.0
2.5
2.0
1.5
1.0
0.5
0.0


Figure S9

a

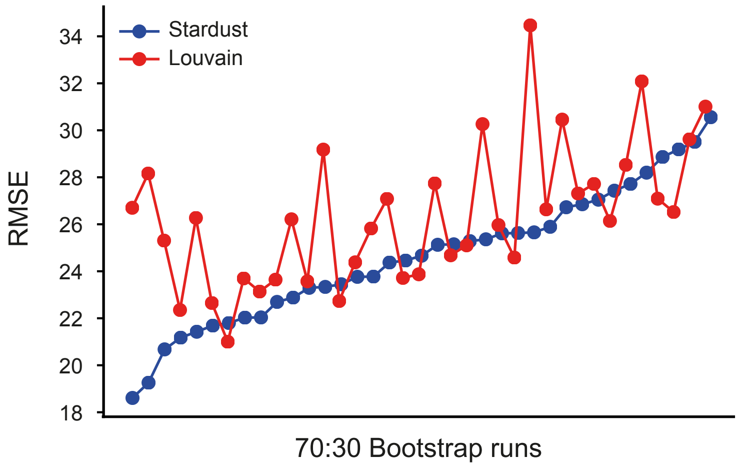


b

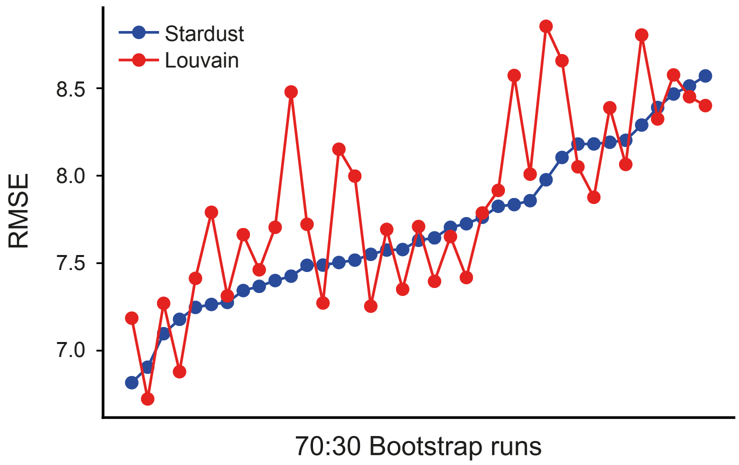


c

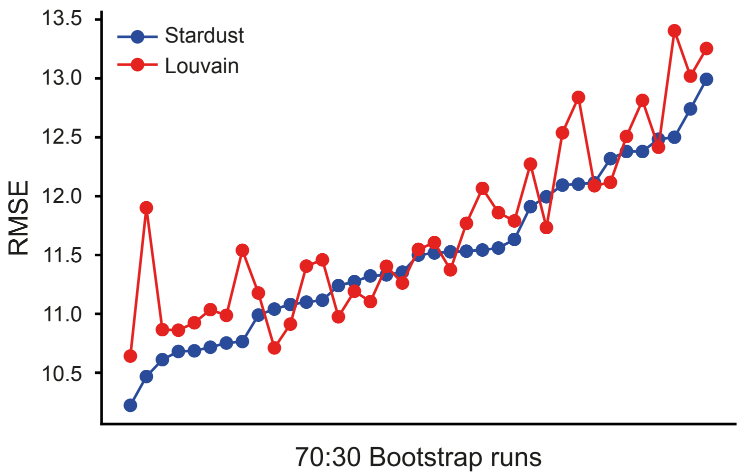


Figure S10

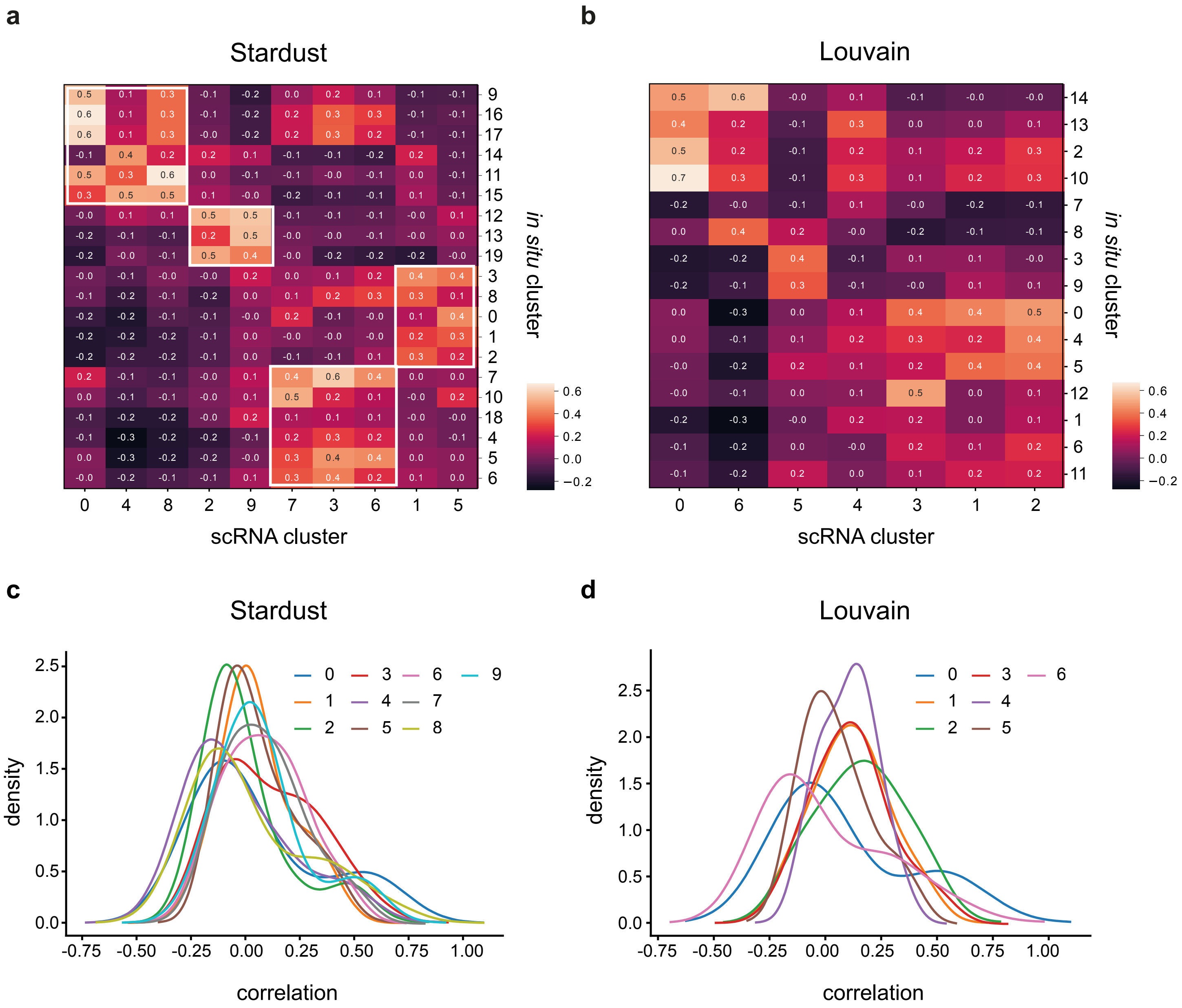
a
Stardust
scRNA cluster
in situ cluster
0 4 8 2 9 7 3 6 1 5
9 16 17 14 11 15 12 13 19 3 8 0 1 2 7 10 18 4 5 6
0.6 0.4 0.2 0.0 −0.2
b
Louvain
scRNA cluster
in situ cluster
0 6 5 4 3 1 2
14 13 2 10 7 8 3 9 0 4 5 12 1 6 11
0.6 0.4 0.2 0.0 −0.2
c
Stardust
density
correlation
0 1 2 3 4 5 6 7 8 9
d
Louvain
density
correlation
0 1 2 3 4 5 6

Figure S11
